# Distinct Microbes, Metabolites, and Ecologies Define the Microbiome in Deficient and Proficient Mismatch Repair Colorectal Cancers

**DOI:** 10.1101/346510

**Authors:** Vanessa L. Hale, Patricio Jeraldo, Jun Chen, Michael Mundy, Janet Yao, Sambhawa Priya, Gary Keeney, Kelly Lyke, Jason Ridlon, Bryan A. White, Amy J. French, Stephen N. Thibodeau, Christian Diener, Osbaldo Resendis-Antonio, Jaime Gransee, Tumpa Dutta, Xuan-Mai Petterson, Ran Blekhman, Lisa Boardman, David Larson, Heidi Nelson, Nicholas Chia

## Abstract

**Background**

The link between colorectal cancer (CRC) and the gut microbiome has been established, but the specific microbial species and their role in carcinogenesis remain controversial. Our understanding would be enhanced by better accounting for tumor subtype, microbial community interactions, metabolism, and ecology.

**Methods**

We collected paired colon tumor and normal–adjacent tissue and mucosa samples from 83 individuals who underwent partial or total colectomies for CRC. Mismatch repair (MMR) status was determined in each tumor sample and classified as either deficient MMR (dMMR) or proficient MMR (pMMR) tumor subtypes. Samples underwent 16S rRNA gene sequencing and a subset of samples from 50 individuals were submitted for targeted metabolomic analysis to quantify amino acids and short-chain fatty acids. A PERMANOVA was used to identify the biological variables that explained variance within the microbial communities. dMMR and pMMR microbial communities were then analyzed separately using a generalized linear mixed effects model that accounted for MMR status, sample location, intra–subject sample correlation, and read depth. Genome–scale metabolic models were then used to generate microbial interaction networks for dMMR and pMMR microbial communities. We assessed global network properties as well as the metabolic influence of each microbe within the dMMR and pMMR networks.

**Results**

We demonstrate distinct roles for microbes in dMMR and pMMR CRC. Sulfidogenic *Fusobacterium nucleatum* and hydrogen sulfide production were significantly enriched in dMMR CRC, but not pMMR CRC. We also surveyed the butyrate–producing microbial species, but did not find a significant difference in predicted or actual butyrate production between dMMR and pMMR microbial communities. Finally, we observed that dMMR microbial communities were predicted to be less stable than pMMR microbial communities. Community stability may play an important role in CRC development, progression, or immune activation within the respective MMR subtypes.

**Conclusions**

Integrating tumor biology and microbial ecology highlighted distinct microbial, metabolic, and ecological properties unique to dMMR and pMMR CRC. This approach could critically improve our ability to define, predict, prevent, and treat colorectal cancers.

## Introduction

The gut microbiota has been linked to colorectal cancer (CRC) in many studies [1–7], and serves as a very promising target for diagnostic, prophylactic, and therapeutic applications. Yet, despite intense study, only a few microbial species—like *Fusobacterium* species—are consistently observed across studies [5–8], while many microbial associations appear to be cohort–specific. Meta–analyses have attempted to overcome the limited statistical power of smaller studies [9] but are limited by the strong biases introduced through varying collection, sequencing, and data processing methodologies [10–13]. Mechanistic studies in mouse models have identified strong causative links between specific microbes (e.g. *Fusobacterium nucleatum, Bacteroides fragilis*) and CRC development and progression [14–17], but these models have limited applicability in genetically diverse human populations. Capturing some of this genetic diversity, on the other hand, may improve our ability to discriminate tumor and normal microbial communities and more clearly define pathways to CRC.

There are multiple subtypes of CRC: one broad categorization is based on mismatch repair (MMR) status. MMR status divides CRCs into two groups: deficient mismatch repair (dMMR) and proficient mismatch repair (pMMR)[18]. In general, dMMR CRCs are hypermethylated, hypermutated, and associated with BRAF V600E mutations; whereas, pMMR CRCs are generally microsatellite stable (MSS) and associated with KRAS[19]. These distinct molecular subtypes of CRC are also borne out by evidence that dMMR is specifically associated with the β-catenin signaling pathway[20]. Clinically, MMR status is associated with patient prognosis, and age, as well as tumor location and stage: Specifically, dMMR CRCs have a better prognosis and occur more often on the right side of the colon in older patients with early stage CRC[18].

Finally, dMMR and pMMR CRC not only have different endpoints, but may also have different paths to tumorigenesis[21] as supported emerging evidence that dMMR CRC arises from sessile serrated adenomas [22] as opposed to the more classic tubular adenoma associated with pMMR CRC [22].

The distinct phenotype of dMMR CRC suggests that host—and possibly also microbial—dynamics are greatly altered in association with deficient mismatch repair. However few CRC microbiome studies account for MMR status [23, 24] or microbial dynamics [2], and no studies, to our knowledge, have assessed both MMR status and microbial community dynamics.

Here, we undertook a new approach in a study involving 83 patients who underwent partial or total colectomy for CRC. From each patient, we collected colon tissue and mucosal samples at tumor and normal–adjacent sites. MMR status was extracted from patient records or determined by testing formalin–fixed paraffin embedded tumor tissue for the expression of four MMR proteins (MLH1, MSH2, MSH6, PMS2). Patient tumors were characterized as either deficient (dMMR) or proficient (pMMR) mismatch repair. Microbial composition was assessed via 16S rRNA gene sequencing. A subset of colon tissue samples additionally underwent metabolomic analysis to quantify amino acids and short-chain fatty acids (SCFAs). A portion of these data was published previously [2] in a study that highlighted the value of integrating *in silico* genome–scale metabolic model predictions and *in vivo* experimental metabolomic data.

From these data, we assessed the relative importance of MMR status compared to other biological factors reported to alter the microbiome[25]. MMR status was the strongest predictor of microbial community variance in comparison to sample location (proximal/distal and on/off tumor), body mass index (BMI), age, and sex. Separate analyses of the dMMR and pMMR microbial communities revealed that many common CRC–associated microbial signatures[9, 26]—including *Fusobacterium nucleatum, Fusobacterium periodonticum,* and *Bacteroides fragilis—*were all enriched in dMMR but not pMMR tumors. Functional differences were examined using a combination of metabolomics and community metabolic modeling. Our results indicate greater predicted and actual hydrogen sulfide production in dMMR CRC as compared to pMMR CRC, but no significant differences in predicted or actual butyrate production. Finally, we approximated microbial ecology by modeling the metabolic interactions between microbes. Overall, the pMMR microbial network was predicted to be more stable (resistant to disturbances). Microbial community stability may play an important role in tumorigenesis, cancer progression, and immune activation [22, 27, 28], and only by examining predicted microbial community interactions were we able to capture this dynamic. Our work demonstrates distinct microbial, metabolic, and ecological attributes of dMMR and pMMR microbial communities, serving to further emphasize the importance of considering tumor biology and microbial interactions in studies of the CRC microbiome.

## Methods

### Human subject enrollment

This study was performed with the approval of the Mayo Clinic Institutional Review Board (IRB# 14-007237 and IRB# 622-00). Written informed consent was obtained from all individuals in the study. Adults (older than 18 years old) who were determined to be candidates for colorectal cancer surgery were voluntarily enrolled at Mayo Clinic in Rochester, Minnesota. Exclusion criteria included chemotherapy or radiation in the 2 weeks leading up to enrollment. Total or partial colectomies were performed on every patient, and colon tissue and mucosal samples were collected from tumor and normal–adjacent sites. Sample location was defined as follows: “proximal” samples were derived from the cecum and ascending colon. “Distal” samples were derived from the transverse, descending, or sigmoid colon, or rectum. MMR status was determined in 83 patients: 25 had dMMR CRC and 58 had pMMR CRC (**Table 1**). We used univariable logistic regression (R v3.1.2) to compare demographic (age, sex, BMI, smoking history) and disease features (tumor location and stage) between dMMR and pMMR groups.

**Table 1.**
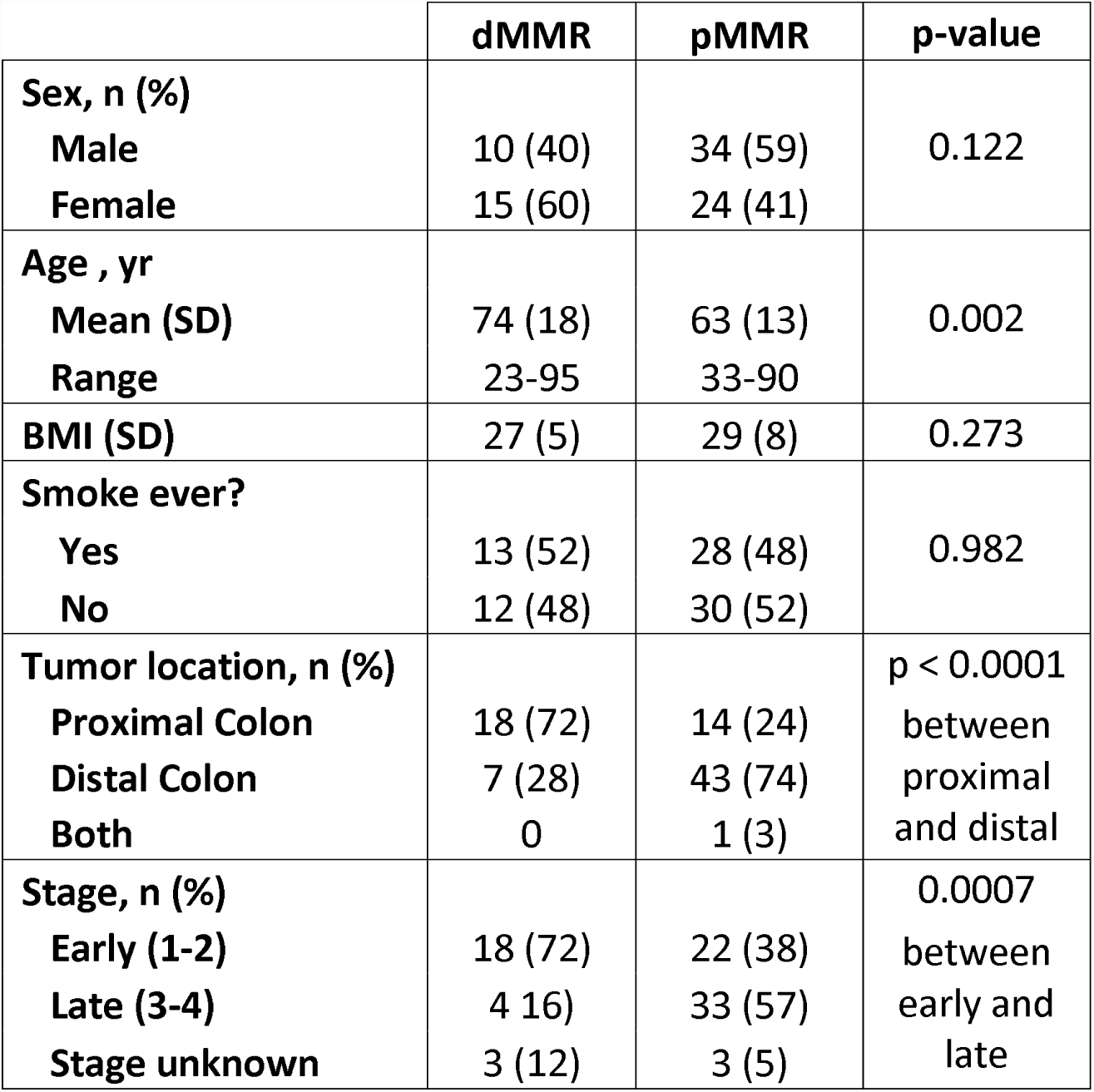
Demographic and disease features of individuals identified as having dMMR or pMMR CRC.

### MMR status determination

Mismatch repair (MMR) pathway and microsatellite instability (MSI) test results were extracted from patient records if available. For patients without MMR test results, banked formalin–fixed paraffin–embedded colon tumor tissue blocks were submitted to the Mayo Clinic Pathology Resource Core for sectioning into 10 micron–thick slices. Slices were then submitted to the Mayo Clinic Molecular Genetics Laboratory for immunohistochemistry staining of MMR proteins (MLH1, PMS2, MSH2, MSH6).

#### 16S DNA extraction, sequencing, and sequence processing

DNA extraction[26] and library preparation on colon tissue (tumor and normal–adjacent), and mucosa were performed as described previously in the Mayo Clinic Microbiome Laboratory [2]. Samples were submitted for 16S rRNA gene sequencing (V3–V5 region) at the Mayo Clinic Medical Genomics Facility (Illumina MiSeq, 2×300, 600 cycles, Illumina Inc.). Sequencing yielded a total of 41,400,384 reads with a median of 70,208 reads per sample. Reads were processed using DADA2 v1.6 to obtain error–corrected amplicon sequence variant representatives—analogous to operational taxonomic units with single-nucleotide resolution (sOTUs) [29]. sOTUs were annotated with genus–level taxonomy using the RDP Naïve Bayesian Classifier[30] as implemented in DADA2 and, if possible, to species level using DADA2, both against the SILVA 16S database, v132[31]. sOTUs annotated as Chloroplast and Mitochondria were removed. Resulting sOTUs were filtered for possible non-specific amplification using SortMeRNA v2.0[32] and Infernal v1.1.2[33]. sOTUs with fewer than 10 reads across all samples were excluded. Multiple sequence alignment of the sOTUs was performed using Infernal v1.1.2[33], and an approximate Maximum Likelihood phylogeny was calculated using FastTree v2.1.9[34]. Raw sequencing data can be found at the NCBI Sequence Read Archive with primary BioProject accession number PRJNA445346 (**Additional File 2** - sOTU table, **3** - sOTU taxonomy, **4**- sOTU fasta file).

### Statistical analyses of 16S rRNA microbial community data

An unweighted UniFrac distance matrix [35] based on the microbial communities in all samples was generated using the phyloseq[36] package v1.22.3. A permutational multivariate analysis of variance (PERMANOVA) was then performed on the distance matrix to assess the effects of MMR status and sample location (proximal/distal and on/off tumor) on variance between microbial communities. The PERMANOVA additionally accounted for subject age, sex, BMI, and sample type (mucosa versus colon tissue) and was performed using the adonis2 function in the vegan[37] package v2.5-1, with 999 bootstrap iterations.

A Generalized Linear Mixed Model (GLMM)[38] was calculated for each sOTU to estimate its abundance (read counts) in relation to predictors that included MMR status and sample location (proximal/distal and on/off tumor). Models were corrected for subject intervariability, specimen type (mucosal vs tissue biopsy), and sequencing read depth, allowing for interactions. We used the package glmmTMB[39] v0.1.4 to estimate the abundance of each microbe under a zero–inflated Poisson distribution. For each predictor, sOTUs were excluded where the method did not converge or the Akaike Information Criterion (AIC) for model quality was not defined. Multiple hypothesis correction was calculated using the Benjamini–Hochberg procedure.

### Validation of differentially abundant microbes using an independent cohort

To validate the differentially abundant microbes associated with dMMR status, we investigated data from a recent study that included microbiome profiling in tumor and matched normal tissue samples in 44 CRC patients [1]. We categorized MMR status based on microsatellite instability (MSI) / microsatellite stable (MSS) status (MSI was categorized as dMMR; MSS was categorized as pMMR) or downregulation of any of the 4 MMR genes (MLH1, MSH2, MSH6 and PMS2) as assessed using RNA-Seq in the same samples. A cutoff (log2(normal/tumor) > = 1) was used to call a gene as downregulated in tumor. Altogether, we identified 9/44 patients as dMMR and the remaining 35/44 as pMMR. Using 16S rRNA gene microbiome characterization for these samples (as described in detail in [1]) we identified sOTUs associated with dMMR tumor/normal and pMMR tumor/normal conditions. We first filtered rare sOTUs, only preserving sOTUs found in at least 50% of our samples, and then performed differential abundance analysis using phyloseq[36] (which uses DESeq2 to build negative binomial generalized linear models). We used the Benjamini–Hochberg method to control for the false discovery rate (FDR).

### Real-time PCR for the *Bacteroides fragilis* toxin gene

Real-time PCR was performed as described previously [2] to test colon tissue and mucosal samples for the presence of the *Bacteroides fragilis* toxin (BFT) genes in 22 dMMR individuals and 53 pMMR individuals. Primers included: BFT-F (5’-GGATAAGCGTACTAAAATACAGCTGGAT-3’), BFT-R (5’-CTGCGAACTCATCTCCCAGTATAAA-3’), and the probe (5’-FAM-CAGACGGACATTCTC-NFQ-MGB-3’)[14].

### Modeling microbial hydrogen sulfide production

We predicted hydrogen sulfide production within dMMR and pMMR tumor and normal–associated microbial communities as described previously[2]. Briefly, we aligned 16S rRNA gene sequences for dMMR tumor and normal samples (colon tissue and mucosa) and pMMR tumor and normal samples against complete genomes in PATRIC and then generated genome–scale metabolic models of each microbe (**Additional File 1: Table S1**). Genome–scale metabolic models use gene annotations from a microbial genome to predict the metabolic inputs and outputs of that microbe. To predict how a microbe might interact within a community, we used MICOM, an open-source platform to assess microbial metabolic community interactions <u>(https://github.com/resendislab/micom).</u> Specifically, we evaluated hydrogen sulfide flux as a measure of hydrogen sulfide production within each microbial community.

### Metabolomics sample preparation and analysis

Colon tissue and mucosa samples were prepared and run as described previously[2]. In brief, UPLC–MS was used to quantify amino acid proxies for hydrogen sulfide including serine, homoserine, lanthionine, L-cystathionine, and D-cystathionine. GC–MS was used to quantify SCFAs including acetate, propionate, isobutyrate, butyrate, isovaleric acid, valeric acid, isocaproic acid, and hexanoate[2]. Significance testing was performed using Kruskal–Wallis and Dunn’s *post hoc* tests in R v3.4.1

### Microbial influence network

To select sOTUs for the Microbial Influence Networks (MINs), we used GLMM results to choose tumor and normal–associated microbes in dMMR and pMMR samples with a linear effect size greater than 0.25, regardless of statistical significance. Effect size captures biological impact potential while significance measures certainty. In this case, we wanted to assess the metabolic influence (i.e., biological impact) of microbes in relation to their respective microbial communities; as such, it was more appropriate to filter by effect size. For each sOTU, the 16S rRNA gene consensus sequence was aligned against complete genome in the PATRIC system using VSEARCH v2.7.1, with a minimum nucleotide identity of 90%. When this procedure generated multiple top hits, we selected a genome, in order, to the most complete genome (fewer contigs), a type strain, a strain with a binomial name, and the closest match to the 16S taxonomy (when possible). For each genome, we then reconstructed and downloaded its corresponding genome–scale metabolic model using the PATRIC service. When sOTUs mapped to the same model, we used that model only once, effectively merging those sOTUs in further analysis, with an exception for when two sOTUs were associated with opposite conditions (i.e., tumor and normal–adjacent samples), in which case we discarded that model from further consideration. The decision to discard was also based on the observation that low identity hits or sOTUs with taxonomy not sufficiently resolved were typically involved in these few cases.

After obtaining the genome–scale metabolic models (GEMs), we calculated “growth” on complete media with no oxygen. This was done by calculating optimal metabolic reaction fluxes using a Flux Balance Analysis[40], in which “growth” is the calculated flux of the reaction defining biomass for a microbe. We did this using a tool for assessing microbial metabolic interactions (MMinte) which evaluates the growth of microbes alone and when paired with another microbe [41]. Once single and paired growth values were calculated using the objective function given by MMinte[41], these values were then used to calculate the influence score. The interaction score, α*_ij_*, for each species, *m*, with a different species *x* was calculated as
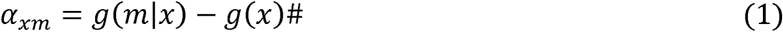

where *g*(*x*) was the growth rate of alone, and *g*(*m*|*x*) was the growth rate of *x* in a community composed of both *x* and *m*. Based on these scores, we then calculated the unweighted metabolic influence of each individual microbial model on the other microbes in the community as the sum of the absolute value of the difference in growth rates when paired with species *m*,
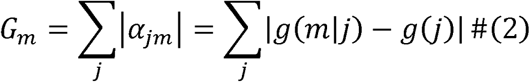

This scoring closely follows the spirit of the scoring from the global interaction modeling in Sung *et al*.[42] with use of actual growth rates instead of summing over shared transporters. Metabolic modeling based on flux balance analysis, as described here, provides a means to calculate a rate in the change of growth, as normalized per unit mass, allowing us to take a simple sum in order to calculate influence under anaerobic conditions.

The percentage of negative interactions was calculated by counting the number of negative interactions over the number of total interactions in each microbial influence network (MIN). Statistical significance was based on the probability of getting equivalent results in dMMR and pMMR networks using the measured distributions of negative and positive interactions in each network and a scheme of random selection with replacement.

Finally, the resulting MIN[42] was visualized using Cytoscape v3.6.1[43] with node and edge properties weighed according to influence score and influence, respectively. Initial visualization in Cytoscape was generated using the “Edge–weighted spring–embedded layout” with the parameters modified to avoid node collisions according to what worked best in each of the two cases. Node sizes and edge weights were likewise set according to the maximum and minimum values in dMMR and pMMR networks separately. Interactions below 10 in the case of dMMR and below 5 in the case of pMMR were excluded from the spring force layout computation in order to achieve better readability of the final network figure. Unconnected nodes that had no influence were not included in the visualization.

### Predicting butyrate production

After we identified the most influential microbes in the dMMR and pMMR MINs, we then assessed the butyrate–producing potential of these microbes. We used the previously selected genome sequences for these microbes and queried the PATRIC service [44] for the presence of the following genes: butyrate kinase (EC 2.7.2.7) or acetate CoA-transferase (EC 2.8.3.8). These genes serve as markers for butyrate production pathways [45]. The presence of either gene was considered sufficient to establish that a bacterium was capable of producing butyrate.

## Results

### dMMR tumors associated with older age and early stage, proximal tumors

A total of 25 individuals with dMMR CRC and 58 individuals with pMMR CRC were involved in this study. Individuals with dMMR CRC were significantly older than individuals with pMMR CRC and significantly more likely to have an early stage, proximal tumor (**Table 1**). As such, we included age and sample location (proximal/distal and on/off tumor) as covariates in subsequent analyses.

### Tumor MMR status strongly predicts variance between microbial communities

To assess factors that contributed to variance in the microbial community data, we performed a PERMANOVA analysis on unweighted UniFrac distances between microbial communities in each sample. We included MMR status, sample location (proximal/distal, on/off tumor), age, sex, BMI and sample type (colon tissue vs. mucosa) as potential predictors of the variance. Remarkably, we found that MMR status explained more of the variance than any of the other 6 variables (**Additional File 1: Table S2**) even when MMR status was included as the last variable in the model (**Table 2**).

**Table 2.** Factors contributing to variance between microbial communities. MMR status was included as the last variable in this model and accounts for the greatest percent variance (PERMANOVA; see also **Additional File 1: Table S2**).

| Factors | % Variation | R<sup>2</sup> | F | Pr(>F) |
| --- | --- | --- | --- | --- |
| BMI | 1.53 | 0.013438 | 8.028927 | 1.00E-04 |
| Age | 1.18 | 0.010349 | 6.183355 | 1.00E-04 |
| Sex | 1.77 | 0.015529 | 9.278172 | 1.00E-04 |
| Sample type | 1.92 | 0.016848 | 10.06614 | 1.00E-04 |
| Sample location - proximal / distal | 2.07 | 0.018191 | 10.86858 | 1.00E-04 |
| Sample location - on/off tumor | 1.26 | 0.011050 | 6.60250 | 1.00E-04 |
| MMR status | 2.11 | 0.018527 | 11.06941 | 1.00E-04 |
| Sample location – proximal / distal:<br>Sample location - on/off tumor | 0.21 | 0.001802 | 1.076725 | 0.3185 |
| Sample location – proximal / distal:MMR<br>status | 1.10 | 0.009634 | 5.7563 | 1.00E-04 |
| Sample location - on/off tumor:MMR<br>status | 0.29 | 0.002572 | 1.537264 | 0.0411 |
| Sample location – proximal / distal:Sample<br>location - on/off tumor:MMR status | 0.19 | 0.001670 | 0.99816721 | 0.4245 |
| Residual | 100.21 | 0.880385 | NA | NA |
| Total | 113.82 | 1 | NA | NA |

### Distinct microbial communities associated with pMMR and dMMR tumors

Given the importance of MMR status to microbial community variance, we opted to assess microbial abundances in tumor and normal samples for each MMR subtype independently. We identified multiple differentially abundant sOTUs in dMMR and pMMR tumor samples as compared to normal–adjacent samples using a generalized linear mixed model (GLMM) that accounted for sample location, sample type, and intrasubject sample correlation (**Fig. 2**; **Additional File 1: Figure S1** (Venn diagram showing counts of microbes in each group**), Table S3** (table of microbes enriched in dMMR), **Table S4** (table of microbes enriched in pMMR). Several major butyrate producers were identified in dMMR and pMMR tumor and normal samples. Only one microbe—*Dorea longicatena*—was significantly enriched in both dMMR and pMMR tumor samples. Four microbes had opposite associations with tumor or normal samples depending on MMR status: *Faecalibacterium prausnitzii* A2-165 and *Blautia* sp. Marseille-P2398 were significantly enriched in pMMR tumor and dMMR normal samples; *Coprococcus comes* ATCC 27758 and *Bacteroides massiliensis* B84634 were significantly enriched in dMMR tumor and pMMR normal samples. Notably, *Fusobacterium periodonticum, F. nucleatum,* and *Bacteroides fragilis*—microbes commonly associated with CRC[5, 14, 46–49]—were among the top most differentially abundant microbes in dMMR tumor samples but were not found to be differentially abundant in pMMR tumor samples.

To validate these results, we used publicly available data from tumor and matched normal samples from 44 CRC patients[1]. Our validation analysis showed several overlapping associations of microbial genomes with respect to dMMR and pMMR in tumor and matched normal samples (**Additional File 1: Table S5, S6)**. dMMR tumors were found enriched for *Bacteroides fragilis* (p=0.02, FDR p=0.37) and *Fusobacterium* (p=0.03, FDR p=0.37) while dMMR normal samples were enriched for *Dorea* (p=0.03, FDR p=0.37) and an Erysipelotrichaceae bacterium (p=0.007, FDR p=0.31) (**Additional File 1: Figure S2**). Even though these associations were not statistically significant after correcting for FDR, their trend of association overlaps with the results from the present study. Differentially abundant sOTUs between pMMR tumors versus normal included Ruminococcaceae, *Faecalibacterium prausnitzii* and *Bacteroides caccae*, which were also differentially abundant in the present study.

As *B. fragilis* was significantly enriched in dMMR tumors, and there are well-established links between toxigenic *B. fragilis* and colorectal cancer [14, 46, 50], we next looked for the presence of the *B. fragilis* toxin (BFT) gene in dMMR and pMMR tissue and mucosa samples. Of the 22 individuals with dMMR CRC, only samples from 1 was BFT positive (5%); of 53 individuals with pMMR CRC, samples from 5 were BFT positive (9.4%). There was no significant difference in BFT presence between individuals with dMMR or pMMR CRC (Chi-squared, p = 0.477).

### Microbial hydrogen sulfide production enriched in the dMMR CRC tumors

As sulfidogenic *F. nucleatum* and *F. periodonticum* were also significantly enriched in dMMR tumor samples, we decided to assess potential hydrogen sulfide production across groups (dMMR/pMMR, tumor/normal) by modeling hydrogen sulfide flux. We used microbial community metabolic models to predict hydrogen sulfide flux within each microbial community (dMMR tumor and normal, pMMR tumor and normal). The models produced a non–significant trend towards increased hydrogen sulfide flux in dMMR tumor samples (**Fig. 3a**). To get a more concrete measure of hydrogen sulfide production, we ran targeted metabolomics to quantify amino acid proxies (serine, homoserine, lanthionine, L-cystathionine, D-cystathionine) for hydrogen sulfide in dMMR and pMMR tumor and normal tissue samples (**Fig. 3b**). We observed a significant increase in lanthionine in dMMR tumor tissue over dMMR or pMMR normal tissue and pMMR tumor. Homoserine and L-Cystathionine were also significantly increased in both dMMR and pMMR tumor tissue as compared to normal–adjacent tissue. The metabolomics results suggest increased hydrogen sulfide production in tumor tissue—particularly in dMMR tumor tissue.

### pMMR microbial community predicted to be more stable and to suppress *F. nucleatum*

To further assess the potential metabolic interactions between tumor and normal–adjacent microbes in relation to MMR status, we constructed two metabolic influence networks (MIN; **Fig. 1**)[42]. The MIN highlights each microbe’s predicted influence and interactions (growth enhancing or suppressing) in relation to other microbes in the community. We also evaluated an indirect measure of ecological stability[51] based on the percentage of predicted positive and negative interactions present within a given microbial community. Notably, the more negative interactions present within a community, the more stable that community is predicted to be[51]. The dMMR microbial community exhibited 21.1% negative interactions while the pMMR microbial community exhibited significantly more (47.6%, binomial test, p<0.0001), suggesting that the pMMR community is more stable.

**Fig. 1.**
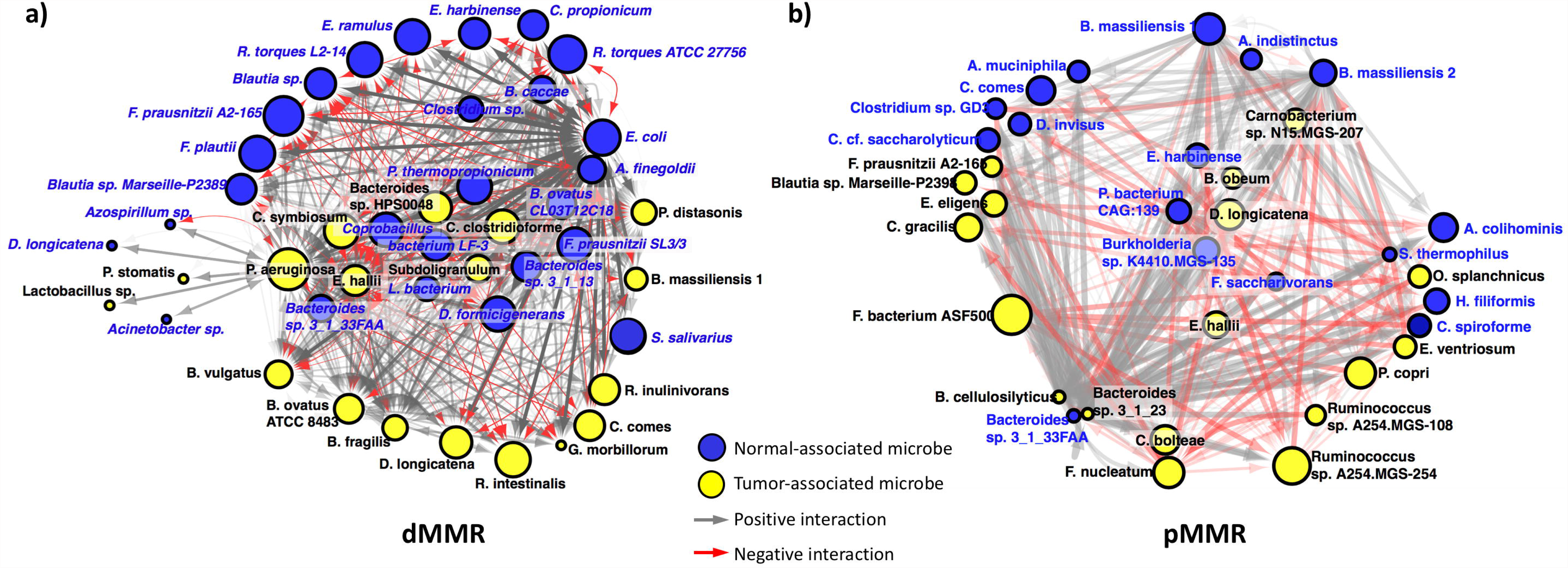
Microbial influence networks for a) dMMR and b) pMMR microbial communities. Node size indicates a microbe’s metabolic influence over other microbes. Edges are directional and weighted and indicate how one microbe affects the growth rate of another: grey edges indicate a positive interaction, i.e., predicted increase in growth when paired, while red edges indicate a negative interaction or a predicted suppression in growth when paired.

**Fig. 2.**
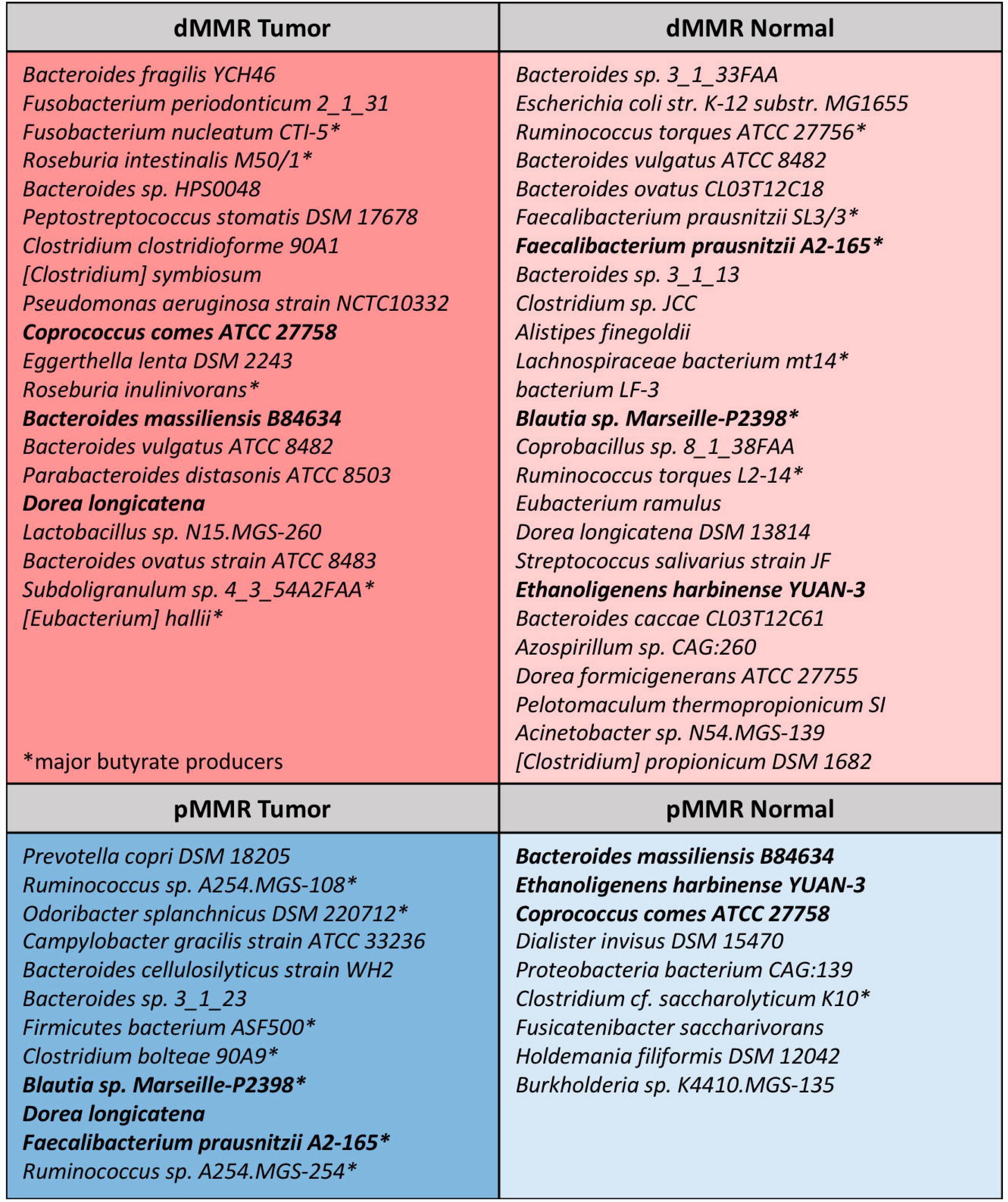
Microbes identified as differentially abundant in tumor as compared to normal samples (tissue and mucosa) from individuals with dMMR or pMMR CRC. Microbes are listed in order of significance from greatest to least. Microbes in bold font are enriched in both dMMR and pMMR samples. For example, *Coprococcus comes* ATCC 27758 is significantly enriched in dMMR tumor samples and pMMR normal samples. (GLMM, all microbes listed have a Benjamini–Hochberg p-value<0.05.)

**Fig. 3.**
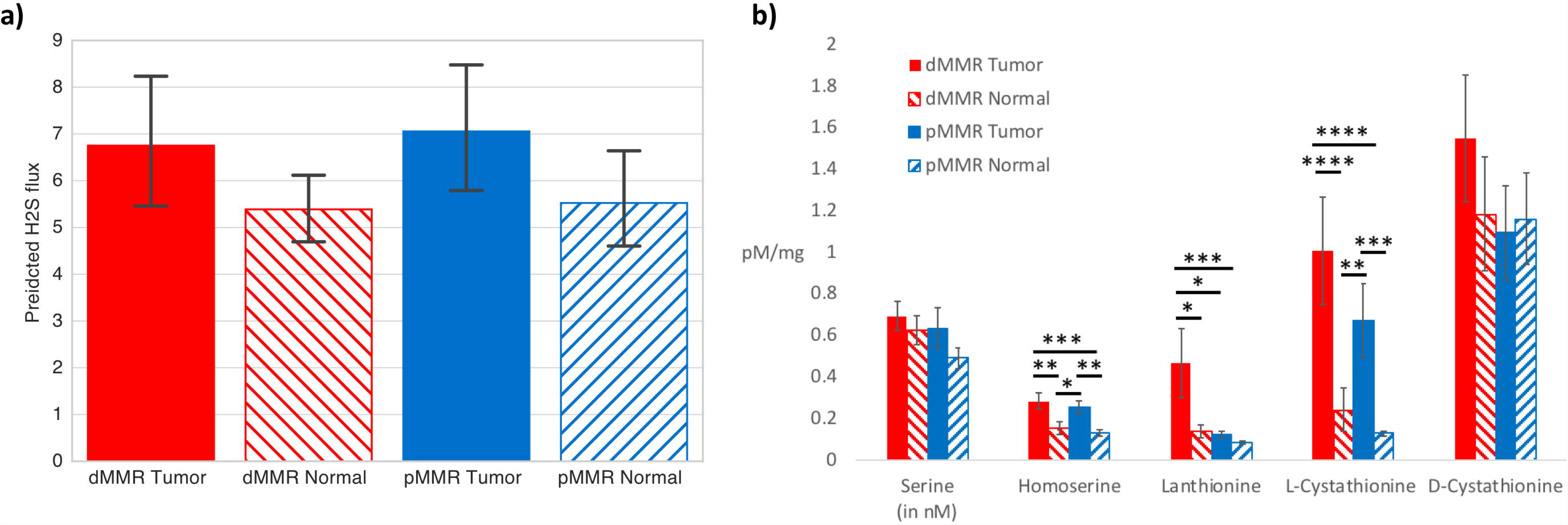
a) Hydrogen sulfide flux predicted based on community metabolic modeling. b) Amino acid proxies for hydrogen sulfide were quantified using UPLC–MS on dMMR and pMMR tumor and normal–adjacent colon tissue samples (Kruskal–Wallis followed by Dunn’s Test for *post hoc* comparisons: *p <0.05; **p<0.0005, ***p<0.0005, ****p<0.00005).

**Fig. 4.**
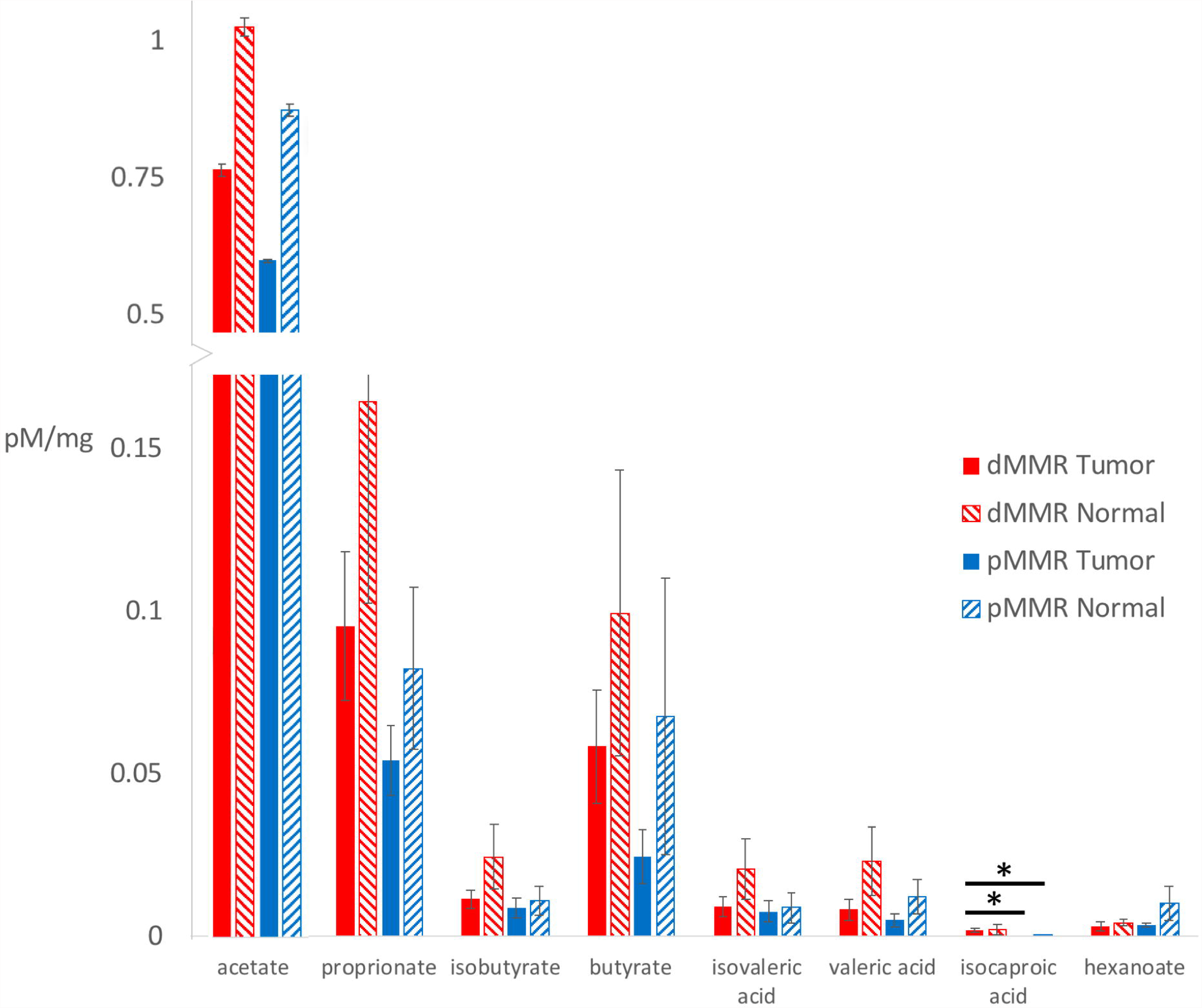
SCFAs in dMMR and pMMR tumor and normal–adjacent colon tissue samples (Kruskal–Wallis followed by Dunn’s Test for *post hoc* comparisons: *p <0.05).

Also of note in relation to the dMMR MIN, *F. nucleatum* and *F. periodonticum* exhibit no metabolic interactions with the other microbes in the network and therefore were not included in the network visualization. In contrast, in the pMMR MIN, *F. nucleatum*—one of the most influential microbes—was uniformly suppressed by 34 out of 44 other pMMR–associated microbes, with zero positive interactions.

### Highly influential microbes include many butyrate producers

Within the MINs, microbes that exhibit many or strong interactions with other microbes—either influencing or being influenced by—are classified as influential microbes. To examine the most influential microbes within each MIN, we identified all microbes with an influence score of 0.5 standard deviations above the mean for dMMR and pMMR microbes, respectively (**Tables 3** – dMMR**, 4 –** pMMR). Characterization of the most influential microbes revealed that many of these microbes are butyrate producers.

**Table 3.** List of basic properties for the most influential microbes in the dMMR MIN.

| dMMR-associated microbes | Influence score | tumor/normal | PATRIC genome ID | butyrate producer? | relevant citations |
| --- | --- | --- | --- | --- | --- |
| *Pseudomonas aeruginosa* | 1732.80 | tumor | 287.2537 |  | [52] |
| *Faecalibacterium prausnitzii* A2-165 | 1615.38 | normal | 411483.3 | x | [53] |
| *Ruminococcus torques* L2-14 | 1516.19 | normal | 657313.3 |  | [54] |
| *Escherichia coli* K-12 | 1452.80 | normal | 511145.12 |  | [55] |
| *Flavonifractor plautii* | 1431.48 | normal | 292800.4 |  | [56] |
| *Eubacterium ramulus* | 1407.40 | normal | 39490.3 |  | [56] |
| *Dorea formicigenerans* | 1407.33 | normal | 411461.4 |  | [57] |
| *Roseburia intestinalis* | 1401.32 | tumor | 657315.3 | x | [53] |
| *Faecalibacterium prausnitzii* SL3/3 | 1395.95 | normal | 657322.3 | x | [53] |
| *Streptococcus salivarius* | 1395.74 | normal | 1304.182 |  | [58] |
| *Ruminococcus torques* ATCC 27756 | 1393.15 | normal | 411460.6 |  | [54] |
| *Clostridium symbiosum* | 1382.77 | tumor | 1512.4 | x | [59] |
| *Pelotomaculum thermopropionicum* | 1376.65 | normal | 370438.4 |  | [60] |
| *Clostridium clostridioforme* | 1368.34 | tumor | 999403.4 | x | [61] |
| *Coprobacillus* sp. 8_1_38FAA | 1363.69 | normal | 450746.3 | x | [62] |

### Butyrate production did not differ between dMMR and pMMR microbial communities

Given the predicted influence and differential abundance of butyrate–producing microbes in dMMR and pMMR, we decided to compare the butyrate–producing potential of the most influential tumor–associated microbes in dMMR and pMMR MINs by searching for genes involved in butyrate production within the functional annotation of these genomes in PATRIC. There were no significant differences in butyrate production–associated genes between dMMR and pMMR tumor–associated microbes (**Table 5;** Wilcoxon rank-sum test, p>0.05). To more fully assess community–wide butyrate production, we performed targeted metabolomics to quantify SCFA concentrations in tumor and normal–adjacent colon tissue. With the exception of isocaproic acid, which was significantly increased in dMMR tumor tissue, there were no significant differences in SCFA concentrations—including butyrate—between dMMR tumor and normal–adjacent or pMMR tumor and normal–adjacent tissue samples (**Fig. 4**).

To further explore how the dMMR MIN could have multiple strongly influential butyrate–producing microbes but no increase in butyrate production, we examined the predicted interactions (positive and negative) of the most influential butyrate producers in the dMMR MIN network (identified in **Table 3**). We found that butyrate producers were targets of more negative interactions (26% negative interactions) as compared to all negative interactions in the dMMR MIN network (21% negative interactions), suggesting growth suppression of butyrate producers in the dMMR community (binomial test, p = 0.02). In contrast, the most influential butyrate producers in the pMMR MIN network (identified in **Table 4**) were not a target for increased negative interactions (binomial test, p = 0.08). We repeated this analysis using all differentially abundant major butyrate producers in dMMR and pMMR microbial communities (identified in **Fig. 2**). We found that while all butyrate producers (in dMMR or pMMR microbial communities) were more likely to be targets of negative interactions, this was more strongly evident in the dMMR microbial community (dMMR binomial test: p=6E-13, pMMR binomial test p=0.008).

**Table 4.** List of basic properties for the most influential microbes in the pMMR MIN.

| pMMR-associated microbes | influence | tumor/normal | PATRIC genome ID | butyrate producer? | relevant citations |
| --- | --- | --- | --- | --- | --- |
| *Firmicutes bacterium ASF500* | 1001.50 | tumor | 1378168.3 |  | [63] |
| *Ruminococcus* sp. A254.MGS-254 | 951.21 | tumor | 1637499.3 |  | [64] |
| *Bacteroides massiliensis* B84634 = | 833.96 | normal | 1121098.3 |  | [65] |
| *Timone 84634 = DSM 17679 = JCM 13223 [PRJNA201686]* |  |  |  |  |  |
| *Fusobacterium nucleatum CTI-5* | 832.56 | tumor | 1316586.3 |  | [47, 66] |
| *Prevotella copri DSM 18205* | 827.75 | tumor | 537011.5 |  | [57] |
| *Dorea longicatena* | 821.96 | tumor | 88431.7 |  | [57] |
| *Clostridium bolteae 90A9* | 797.92 | tumor | 997894.4 | x | [61] |
| *Anaerotruncus colihominis DSM 17241* | 780.61 | normal | 445972.6 | x | [67] |
| *Coprococcus comes ATCC 27758* | 779.84 | normal | 470146.3 | x | [68] |
| *Campylobacter gracilis strain ATCC 33236* | 766.50 | tumor | 824.5 |  | [69] |

**Table 5.** Percent of the most influential tumor–associated microbes in dMMR and pMMR CRC that contain a butyrate–producing pathway, as predicted by the presence of genes involved in butyrate production.

| Gene annotation | EC number | dMMR | pMMR |
| --- | --- | --- | --- |
| Acetate CoA-transferase | 2.8.3.8 | 14.9% | 13.8% |
| Butyrate kinase | 2.7.2.7 | 27.7% | 31.0% |

## Discussion

This study integrates tumor biology and microbial ecology in a novel and powerful approach to understanding colorectal cancer. Our results indicate that MMR status is one of the strongest predictors of microbial community variance; however, few studies [23, 24], to date, include MMR status in microbial community analysis of colorectal cancer. Interestingly, we also identified several differentially abundant microbes associated with dMMR but not pMMR tumor samples including *F. nucleatum, F. periodonticum,* and *B. fragilis.* We further validated these findings in an independent cohort[1], which underscores the importance of including MMR status in future CRC microbiome studies. We additionally characterized the predicted and actual metabolic profiles of dMMR and pMMR individuals in relation to hydrogen sulfide and butyrate production, and we generated a network of predicted interactions within the dMMR and pMMR microbial communities.

Hydrogen sulfide has been reported to both promote and inhibit colorectal cancer [70–73]. To assess the role of hydrogen sulfide within our study, we looked for sulfidogenic bacteria, predicted hydrogen sulfide production using community metabolic models, and measured hydrogen sulfide concentrations through targeted metabolomics for amino acid proxies. We found two significantly enriched hydrogen sulfide–producing *Fusobacterium* species and significantly increased hydrogen sulfide concentrations in dMMR tumor samples. In the microbial influence network, both *Fusobacterium* species exhibited zero predicted interactions—positive or negative—with other microbes in the network. Together, this suggests that these *Fusobacterium* species grow abundantly and unchecked by other microbes, and have the potential to produce large quantities of hydrogen sulfide. In contrast, the *F. nucleatum* found in pMMR MIN was predicted to be the target of 34 (100%) negative interactions, suggesting that its growth and metabolic output (hydrogen sulfide) are highly suppressed in pMMR individuals.

These intriguing results leads us to speculate on the relationship between *Fusobacterium* species, hydrogen sulfide production and dMMR CRC. Notably, *Fusobacterium* species have previously been associated with hypermethylation of MLH1, MSI, BRAF mutations, and poorly differentiated tumors[4, 47]—all of which are characteristics of dMMR CRC[74]. Hydrogen sulfide—a cytotoxic, genotoxic gas—has also been associated with CRC[70, 71], although its role is somewhat controversial[72, 73]. A recent report indicates that colon cancer cells may respond to hydrogen sulfide in a bell–shaped dose–dependent manner: at high concentrations, hydrogen sulfide inhibits the proliferation of cancer cells, while at lower concentrations, hydrogen sulfide can stimulate the proliferation of cancer cells[73, 75]. In dMMR, if high levels of hydrogen sulfide (and hydrogen sulfide producers) inhibit cancer proliferation, then we would expect individuals with dMMR to present with earlier stage cancer—which is indeed the case in our cohort and other reported cohorts [65]. dMMR CRC has also been associated with lower recurrence rates and a better prognosis[74]. In opposition to these findings are studies showing that *F. nucleatum* can potentiate tumorigenesis and that *F. nucleatum*–associated CRCs have a worse prognosis [4, 5].

Besides *Fusobacterium*, *Bacteroides fragilis* was also found to be significantly enriched in dMMR tumor samples. Toxigenic *B. fragilis* has well–established and causative links to inflammation, and CRC[50, 76], and inflammation has been linked to hypermethylation[77]. As such, we tested dMMR and pMMR tissue and mucosa samples for the presence of the *B. fragilis* toxin (BFT) gene but did not find a significant difference in the presence of the BFT gene between dMMR and pMMR individuals. Given these results, it is unclear what the significance of increased *B. fragilis* is in the dMMR tumor samples.

Butyrate has been another subject of intense investigation in relation to gut health and CRC[54, 57]. Conflicting work in murine models indicates that butyrate can act to repress or to accelerate polyp and tumor formation—resulting in the so–called butyrate paradox[78]. Recently, a dMMR CRC mouse model showed that microbially–produced butyrate accelerated tumorigenesis[24], indicating that the source of this paradox may have to do with the genetic model of CRC being used and that butyrate may therefore have different, even opposite, roles in different CRC subtypes. It was therefore intriguing that more butyrate producers were identified as highly influential in the dMMR MIN as compared to pMMR MIN. However, neither predicted (functional annotation) nor actual (metabolomic data) butyrate production differed significantly between dMMR and pMMR samples or tumor and normal samples in the metabolomic profile. One potential reason for this lack of difference surfaced when we examined the number of negative interactions targeted at butyrate producers in the dMMR and pMMR MINs. dMMR butyrate producers had a significantly higher probability of negative interactions as compared to pMMR butyrate producers. This suggests that, though present, these butyrate producers are being suppressed in dMMR microbial communities.

Negative interactions were also predicted to be significantly increased in the pMMR microbial community as a whole versus the dMMR microbial community. Negative interactions, implying competition, are an ecological hallmark of a stable community[51], which is both resistant and resilient to disturbance. We speculate that a less stable microbial community in dMMR individuals could mean constant community shifts and disturbances—resulting in increased immune activation. The dMMR tumor phenotype is associated with increased immune response[74], which may play a role in inhibiting cancer cell proliferation of dMMR tumors.

Overall, our study demonstrates the importance and value in considering tumor biology (MMR status) and ecological interactions when evaluating microbial community data. Our work is primarily descriptive and incorporates host clinical features, microbiome, metabolome, and modeling data. While we make speculations based on these data, future prospective and mechanistic studies are needed to test these ideas. We also recognize that selecting sequenced genomes available in the database to represent 16S rRNA sOTUs cannot fully replace metagenomic sequencing given well–known strain-to-strain variation in gene content. However, these variations between strains are often largely in secondary metabolite pathways, rather than core metabolic function, which is the main target of our modeling analysis.

Another limitation of this study is our inability to attribute a source to metabolomic data. While butyrate is a microbial fermentation product, hydrogen sulfide and its amino acid proxies can be produced by both humans and bacteria. Thus, the enriched hydrogen sulfide we detect in dMMR tumor samples could potentially be attributed to increased hydrogen sulfide production within tumor tissue, and indeed, this has been reported[73]. If this was the solely case here however, we might expect to see similar increases in hydrogen sulfide in pMMR tumors—most of which are later in stage than dMMR tumors. We did not see this, suggesting that it is feasible that the increased hydrogen sulfide production in dMMR tumors is coming from an exogenous (microbial) source. Notably, microbially produced hydrogen sulfide can be generated from multiple pathways including the respiration of dietary taurine and sulfate as well as the degradation of sulfomucins. The amino acid proxies we use to assess hydrogen sulfide production only capture some, but not all of these potential pathways, so we may have underestimated hydrogen sulfide production.

Finally, the field of genome–scale metabolic modeling has only recently encompassed tools for community metabolic analyses[79], and many of the tools[41, 42, 80] are sensitive to the underlying quality of the metabolic models[44, 81]. Models vary greatly depending on the presence and accuracy genome annotations which will generally improve over time. Future work aimed at understanding and verifying microbial dynamics in relation to MMR status or other CRC subtypes could dramatically improve our ability to define, predict, prevent, and treat colorectal cancers.

## Conclusions

This study provides a novel framework in which to examine colorectal cancer:

1. Host–microbe interactions: Tumor MMR status strongly predicted microbial community variance and was associated with distinct microbial, metabolic, and interaction profiles. Our approach incorporating tumor MMR status, microbiome, metabolome and modeling data allowed us unique insights into the role of hydrogen sulfide and hydrogen sulfide producers within the dMMR microbial community. Tumor biology (e.g. MMR status) and microbial ecology are inextricably linked, and it is critical that future studies account for both in order to understand and more precisely classify the many pathways to CRC.
2. Microbe–microbe interactions: Microbial influence networks provided *in silico* predictions of community stability and microbial interactions that aligned with *in vivo* metabolomics data: Suppression of sulfidogenic *F. nucleatum* and significantly lower hydrogen sulfide production in pMMR, and suppression of butyrate–producing bacteria in dMMR—which may explain the lack of difference in butyrate production between dMMR and pMMR samples. The validation of *in silico* data with *in vivo* tests provides support for a future of precision medicine tools that can accurately predict disease and the potential effects of prophylactic or therapeutic interventions on the microbiome.

Microbes act within communities, and understanding and predicting these interactions will be key to developing targeted mechanisms to help prevent or treat colorectal cancer.

## Acknowledgements

We would first like to thank the patients who volunteered for this study. We also thank the many other individuals who made this work possible including members of the Mayo Clinic Microbiome Laboratory, study coordinators, students, colorectal surgeons, program directors, and pathology assistants. We also specially acknowledge Donna Felmlee Devine and Caitlin Foss-Baumgard for their assistance with patient records in relation to this study. Finally, we gratefully acknowledge the following funding sources: NIH (R01CA179243; N.C. and V.L.H. and R01CA170357; L.B.), the Mayo Clinic Center for Cell Signaling in Gastroenterology (NIDDK P30DK084567), the Mayo Clinic Metabolomics Resource Core Pilot and Feasibility Award (U24DK100469), the Fred C. Andersen Foundation (H.N. and N.C.), the Mayo Clinic Center for Individualized Medicine, The Randy Shaver Cancer Research and Community Fund (R.B.) the Minnesota Partnership for Biotechnology and Medical Genomics (R.B.), The Alfred P. Sloan Foundation (R.B.) O.R.A thanks the financial support coming from the National Institute of Genomic Medicine (INMEGEN) to develop the computational tool used for the microbiome analysis (MICOM).

## Additional File 1

**Figure S1:** Venn diagram highlighting number of microbes that overlap between tumor and normal samples in relation to MMR status (dMMR = red font, pMMR = green font, red circles = tumor samples, blue circles = normal samples). Only differentially abundant microbes with a corrected p-value < 0.05 were included in this diagram.

**Figure S2:** Differentially abundant OTUs between patient–matched tumor and normal samples in individuals with dMMR CRC. Y-axis indicates percent relative abundance of OTUs. Line color indicates directionality of change in microbial abundance: red = increased abundance relative to normal, blue = decreased abundance or no change compared to normal.

**Table S1.** Factors contributing to variance between microbial communities. Sample location was included as the last variable in this model (PERMANOVA).

**Table S2.** Factors contributing to variance between microbial communities. Sample location was included as the last variable in this model (PERMANOVA).

**Table S3:** sOTUs enriched in tumor samples (colon tissue and mucosa) as compared to normal–adjacent samples in individuals with dMMR CRC.

**Table S4:** sOTUs enriched in tumor samples (colon tissue and mucosa) as compared to normal–adjacent samples in individuals with pMMR CRC.

**Table S5:** Differentially abundant microbes in individuals with dMMR CRC. Blue boxes highlight microbes enriched in tumor tissue samples as compared to normal–adjacent samples.

**Table S6:** Differentially abundant microbes in individuals with pMMR CRC. Blue boxes highlight microbes enriched in tumor tissue samples as compared to normal–adjacent samples.

## References

1. Burns MB, Montassier E, Abrahante J, Priya S, Niccum DE, Khoruts A, et al. Colorectal cancer mutational profiles correlate with defined microbial communities in the tumor microenvironment. bioRxiv. 2018;:090795. doi:10.1101/090795.

2. Hale VL, Jeraldo P, Mundy M, Yao J, Keeney G, Scott N, et al. Synthesis of multi-omic data and community metabolic models reveals insights into the role of hydrogen sulfide in colon cancer. Methods. 2018. doi:10.1016/j.ymeth.2018.04.024.

3. Flemer B, Lynch DB, Brown JMR, Jeffery IB, Ryan FJ, Claesson MJ, et al. Tumour-associated and non-tumour-associated microbiota in colorectal cancer. Gut. 2017;66:633–43. doi:10.1136/gutjnl-2015-309595.

4. Mima K, Nishihara R, Qian ZR, Cao Y, Sukawa Y, Nowak JA, et al. Fusobacterium nucleatum in colorectal carcinoma tissue and patient prognosis. Gut. 2016;65:1973–80. doi:10.1136/gutjnl-2015-310101.

5. Kostic AD, Chun E, Robertson L, Glickman JN, Gallini CA, Michaud M, et al. Fusobacterium nucleatum Potentiates Intestinal Tumorigenesis and Modulates the Tumor-Immune Microenvironment. Cell Host Microbe. 2013;14:207–15.

6. Chen W, Liu F, Ling Z, Tong X, Xiang C. Human Intestinal Lumen and Mucosa-Associated Microbiota in Patients with Colorectal Cancer. PLoS One. 2012;7:e39743. doi:10.1371/journal.pone.0039743.

7. Flanagan L, Schmid J, Ebert M, Soucek P, Kunicka T, Liska V, et al. Fusobacterium nucleatum associates with stages of colorectal neoplasia development, colorectal cancer and disease outcome. Eur J Clin Microbiol Infect Dis. 2014.

8. Castellarin M, Warren RL, Freeman JD, Dreolini L, Krzywinski M, Strauss J, et al. Fusobacterium nucleatum infection is prevalent in human colorectal carcinoma. Genome Res. 2012;22:299–306. doi:10.1101/gr.126516.111.

9. Drewes JL, White JR, Dejea CM, Fathi P, Iyadorai T, Vadivelu J, et al. High-resolution bacterial 16S rRNA gene profile meta-analysis and biofilm status reveal common colorectal cancer consortia. npj Biofilms Microbiomes. 2017;3.

10. Sinha R, Abu-Ali G, Vogtmann E, Fodor AA, Ren B, Amir A, et al. Assessment of variation in microbial community amplicon sequencing by the Microbiome Quality Control (MBQC) project consortium. Nat Biotechnol. 2017;35:1077.

11. Vogtmann E, Chen J, Amir A, Shi J, Abnet CC, Nelson H, et al. Comparison of collection methods for fecal samples in microbiome Studies. Am J Epidemiol. 2017;185:115–23.

12. Sinha R, Chen J, Amir A, Vogtmann E, Shi J, Inman KS, et al. Collecting Fecal Samples for Microbiome Analyses in Epidemiology Studies. Cancer Epidemiol Biomarkers Prev. 2016;25:407–16. doi:10.1158/1055-9965.EPI-15-0951.

13. Jeraldo P, Chia N, Goldenfeld N. On the suitability of short reads of 16S rRNA for phylogeny-based analyses in environmental surveys. Environ Microbiol. 2011;13:3000–9. doi:10.1111/j.1462-2920.2011.02577.x.

14. Housseau F, Sears CL. Enterotoxigenic Bacteroides fragilis (ETBF)-mediated colitis in Min (Apc+/−) mice: A human commensal-based murine model of colon carcinogenesis. Cell Cycle. 2010;9:3–5.

15. Zackular JP, Baxter NT, Chen GY, Schloss PD. Manipulation of the Gut Microbiota Reveals Role in Colon Tumorigenesis. mSphere. 2016;1:e00001–15. doi:10.1128/mSphere.00001-15.

16. Bishehsari F, Engen PA, Preite NZ, Tuncil YE, Naqib A, Shaikh M, et al. Dietary fiber treatment corrects the composition of gut microbiota, promotes SCFA production, and suppresses colon carcinogenesis. Genes (Basel). 2018;9.

17. Khazaie K, Zadeh M, Khan MW, Bere P, Gounari F, Dennis K, et al. Abating colon cancer polyposis by Lactobacillus acidophilus deficient in lipoteichoic acid. Proc Natl Acad Sci. 2012;109:10462–7.

18. French AJ, Sargent DJ, Burgart LJ, Foster NR, Kabat BF, Goldberg R, et al. Prognostic significance of defective mismatch repair and BRAF V600E in patients with colon cancer. Clin Cancer Res. 2008;14:3408–15.

19. Guinney J, Dienstmann R, Wang X, De Reyniès A, Schlicker A, Soneson C, et al. The consensus molecular subtypes of colorectal cancer. Nat Med. 2015;21:1350.

20. Mårtensson A, Oberg A, Jung A, Cederquist K, Stenling R, Palmqvist R. Beta-catenin expression in relation to genetic instability and prognosis in colorectal cancer. Oncol Rep. 2007;17:447–52.

21. Morkel M, Riemer P, Bläker H, Sers C. Similar but different: distinct roles for KRAS and BRAF oncogenes in colorectal cancer development and therapy resistance. Oncotarget. 2015;6:20785–800.

22. Sweetser S, Jones A, Smyrk TC, Sinicrope FA. Sessile Serrated Polyps are Precursors of Colon Carcinomas With Deficient DNA Mismatch Repair. Clin Gastroenterol Hepatol. 2016;14:1056–9.

23. Purcell R V., Visnovska M, Biggs PJ, Schmeier S, Frizelle FA. Distinct gut microbiome patterns associate with consensus molecular subtypes of colorectal cancer. Sci Rep. 2017;7:11590. doi:10.1038/s41598-017-11237-6.

24. Belcheva A, Irrazabal T, Robertson SJJ, Streutker C, Maughan H, Rubino S, et al. Gut Microbial Metabolism Drives Transformation of Msh2-Deficient Colon Epithelial Cells. Cell. 2014;158:288–99. doi:10.1016/j.cell.2014.04.051.

25. Chen J, Ryu E, Hathcock M, Ballman K, Chia N, Olson JE, et al. Impact of demographics on human gut microbial diversity in a US Midwest population. PeerJ. 2016;4:e1514. doi:10.7717/peerj.1514.

26. Hale VL, Chen J, Johnson S, Harrington SC, Yab TC, Smyrk TC, et al. Shifts in the fecal microbiota associated with adenomatous polyps. Cancer Epidemiol Prev Biomarkers. 2017;26:85–94.

27. Sweetser S, Smyrk TC, Sinicrope FA. Serrated colon polyps as precursors to colorectal cancer. Clinical Gastroenterology and Hepatology. 2013;11:760–7.

28. Rashtak S, Rego R, Sweetser SR, Sinicrope FA. Sessile serrated polyps and colon cancer prevention. Cancer Prevention Research. 2017;10:270–8.

29. Callahan BJ, McMurdie PJ, Rosen MJ, Han AW, Johnson AJA, Holmes SP. DADA2: High-resolution sample inference from Illumina amplicon data. Nat Methods. 2016;13:581–3. doi:10.1038/nmeth.3869.

30. Wang Q, Garrity GM, Tiedje JM, Cole JR. Naive Bayesian classifier for rapid assignment of rRNA sequences into the new bacterial taxonomy. Appl Environ Microbiol. 2007;73:5261–7. doi:10.1128/AEM.00062-07.

31. Quast C, Pruesse E, Yilmaz P, Gerken J, Schweer T, Yarza P, et al. The SILVA ribosomal RNA gene database project: improved data processing and web-based tools. Nucleic Acids Res. 2013;41 Database issue:D590–6. doi:10.1093/nar/gks1219.

32. Kopylova E, Noé L, Touzet H. SortMeRNA: Fast and accurate filtering of ribosomal RNAs in metatranscriptomic data. Bioinformatics. 2012;28:3211–7.

33. Nawrocki EP, Eddy SR. Infernal 1.1: 100-fold faster RNA homology searches. Bioinformatics. 2013;29:2933–5. doi:10.1093/bioinformatics/btt509.

34. Price MN, Dehal PS, Arkin AP. FastTree 2 -- Approximately Maximum-Likelihood Trees for Large Alignments. PLoS One. 2010;5:e9490. doi:10.1371/journal.pone.0009490.

35. Lozupone C, Knight R. UniFrac: a New Phylogenetic Method for Comparing Microbial Communities. Appl Environ Microbiol. 2005;71:8228–35.

36. McMurdie PJ, Holmes S. phyloseq: An R Package for Reproducible Interactive Analysis and Graphics of Microbiome Census Data. PLoS One. 2013;8:e61217. doi:10.1371/journal.pone.0061217.

37. Oksanen J, Kindt R, Legendre P, O’Hara B, Simpson GL, Solymos PM, et al. The vegan package. Community Ecol Packag. 2008;:190.

38. Bolker BM, Brooks ME, Clark CJ, Geange SW, Poulsen JR, Stevens MHH, et al. Generalized linear mixed models: a practical guide for ecology and evolution. Trends Ecol Evol. 2009;24:127–35.

39. Brooks ME, Kristensen K, van Benthem KJ, Magnusson A, Berg CW, Nielsen A, et al. glmmTMB balances speed and flexibility among packages for zero-inflated generalized linear mixed modeling. R J. 2017;9:378–400.

40. Orth JD, Thiele I, Palsson BØ. What is flux balance analysis? Nat Biotechnol. 2010;28:245–8. doi:10.1038/nbt.1614.

41. Mendes-Soares H, Mundy M, Soares LM, Chia N. MMinte: an application for predicting metabolic interactions among the microbial species in a community. BMC Bioinformatics. 2016;17:343.

42. Sung J, Kim S, Cabatbat JJT, Jang S, Jin Y-S, Jung GY, et al. Global metabolic interaction network of the human gut microbiota for context-specific community-scale analysis. Nat Commun. 2017;8:15393.

43. Shannon P, Markiel A, Ozier O, Baliga NS, Wang JT, Ramage D, et al. Cytoscape: A software Environment for integrated models of biomolecular interaction networks. Genome Res. 2003;13:2498–504.

44. Wattam AR, Abraham D, Dalay O, Disz TL, Driscoll T, Gabbard JL, et al. PATRIC, the bacterial bioinformatics database and analysis resource. Nucleic Acids Res. 2014;42:D581–91. doi:10.1093/nar/gkt1099.

45. Anand S, Kaur H, Mande SS. Comparative In silico Analysis of Butyrate Production Pathways in Gut Commensals and Pathogens. Front Microbiol. 2016;7:1945. doi:10.3389/fmicb.2016.01945.

46. Chung L, Thiele Orberg E, Geis AL, Chan JL, Fu K, DeStefano Shields CE, et al. Bacteroides fragilis Toxin Coordinates a Pro-carcinogenic Inflammatory Cascade via Targeting of Colonic Epithelial Cells. Cell Host Microbe. 2018;23:203–214.e5.

47. Koi M, Okita Y, M. Carethers J. Fusobacterium nucleatum Infection in Colorectal Cancer: Linking Inflammation, DNA Mismatch Repair and Genetic and Epigenetic Alterations. J Anus, Rectum Colon. 2018;2:37–46.

48. Kostic AD, Gevers D, Pedamallu CS, Michaud M, Duke F, Earl AM, et al. Genomic analysis identifies association of Fusobacterium with colorectal carcinoma. Genome Res. 2012;22:292–8. doi:10.1101/gr.126573.111.

49. Abed J, Emgård JEM, Zamir G, Faroja M, Almogy G, Grenov A, et al. Fap2 Mediates Fusobacterium nucleatum Colorectal Adenocarcinoma Enrichment by Binding to Tumor-Expressed Gal-GalNAc. Cell Host Microbe. 2016;20:215–25. doi:10.1016/j.chom.2016.07.006.

50. Dejea CM, Fathi P, Craig JM, Boleij A, Taddese R, Geis AL, et al. Patients with familial adenomatous polyposis harbor colonic biofilms containing tumorigenic bacteria. Science (80-). 2018;359:592–7. doi:10.1126/science.aah3648.

51. Coyte KZ, Schluter J, Foster KR. The ecology of the microbiome: Networks, competition, and stability. Science (80-). 2015;350:663–6. doi:10.1126/science.aad2602.

52. Eschbach M, Schreiber K, Trunk K, Buer J, Jahn D, Schobert M. Long-term anaerobic survival of the opportunistic pathogen Pseudomonas aeruginosa via pyruvate fermentation. J Bacteriol. 2004;186:4596–604.

53. Rivière A, Selak M, Lantin D, Leroy F, De Vuyst L. Bifidobacteria and butyrate-producing colon bacteria: Importance and strategies for their stimulation in the human gut. Frontiers in Microbiology. 2016;7 JUN.

54. Pryde SE, Duncan SH, Hold GL, Stewart CS, Flint HJ. The microbiology of butyrate formation in the human colon. FEMS Microbiology Letters. 2002;217:133–9.

55. Sousa CP. The versatile strategies of Escherichia coli pathotypes: a mini review. Rev Lit Arts Am. 2006;12:363–73.

56. Ulbrich K, Reichardt N, Braune A, Kroh LW, Blaut M, Rohn S. The microbial degradation of onion flavonol glucosides and their roasting products by the human gut bacteria *Eubacterium ramulus* and *Flavonifractor plautii*. Food Res Int. 2015;67:349–55.

57. Louis P, Flint HJ. Formation of propionate and butyrate by the human colonic microbiota. Environmental Microbiology. 2017;19:29–41.

58. Jia F. Genome Sequence of the Oral Probiotic *Streptococcus salivarius* JF. Genome Announc. 2016;4:e00971–16.

59. Van Den Abbeele P, Belzer C, Goossens M, Kleerebezem M, De Vos WM, Thas O, et al. Butyrate-producing Clostridium cluster XIVa species specifically colonize mucins in an in vitro gut model. ISME J. 2013;7:949–61.

60. Imachi H, Sekiguchi Y, Kamagata Y, Hanada S, Ohashi A, Harada H. Pelotomaculum thermopropionicum gen. nov., sp. nov., an anaerobic, thermophilic, syntrophic propionate-oxidizing bacterium. Int J Syst Evol Microbiol. 2002;52:1729–35.

61. Dehoux P, Marvaud JC, Abouelleil A, Earl AM, Lambert T, Dauga C. Comparative genomics of Clostridium bolteae and Clostridium clostridioforme reveals species-specific genomic properties and numerous putative antibiotic resistance determinants. BMC Genomics. 2016;17.

62. Kageyama A, Benno Y. Coprobacillus catenaformis gen. nov., sp. nov., a new genus and species isolated from human feces. Microbiol Immunol. 2000;44:23–8.

63. Wannemuehler MJ, Overstreet A, Ward D V, Phillips GJ. Draft genome sequences of the altered schaedler flora, a defined bacterial community from gnotobiotic mice. Genome Announc. 2014;2:1–2.

64. Jeraldo P, Hernández Á, White BA, O’Brien D, Ahlquist D, Boardman L, et al. Draft Genome Sequences of 24 Microbial Strains Assembled from Direct Sequencing from 4 Stool Samples. Genome Announc. 2015;3:e00526–15. doi:10.1128/genomeA.00526-15.

65. Tatusova T, Ciufo S, Fedorov B, O’Neill K, Tolstoy I. RefSeq microbial genomes database: new representation and annotation strategy (vol 42, pg 553, 2014). Nucleic Acids Res. 2015;43:3872.

66. Han YW. Fusobacterium nucleatum: A commensal-turned pathogen. Current Opinion in Microbiology. 2015.

67. Lawson PA, Song Y, Liu C, Molitoris DR, Vaisanen ML, Collins MD, et al. Anaerotruncus colihominis gen. nov., sp. nov., from human faeces. Int J Syst Evol Microbiol. 2004;54:413–7.

68. Louis P, Flint HJ. Diversity, metabolism and microbial ecology of butyrate-producing bacteria from the human large intestine. FEMS Microbiol Lett. 2009;294:1–8.

69. Shinha T. Fatal Bacteremia Caused by Campylobacter gracilis, United States. Emerg Infect Dis. 2015;21:1084–5. doi:10.3201/eid2106.142043.

70. Attene-Ramos MS, Nava GM, Muellner MG, Wagner ED, Plewa MJ, Gaskins HR. DNA damage and toxicogenomic analyses of hydrogen sulfide in human intestinal epithelial FHs 74 int cells. Environ Mol Mutagen. 2010;51:304–14.

71. Wolf PG, Parthasarathy G, Chen J, O’Connor HM, Chia N, Bharucha AE, et al. Assessing the colonic microbiome, hydrogenogenic and hydrogenotrophic genes, transit and breath methane in constipation. Neurogastroenterol Motil. 2017.

72. Lee ZW, Zhou J, Chen CS, Zhao Y, Tan CH, Li L, et al. The slow-releasing Hydrogen Sulfide donor, GYY4137, exhibits novel anti-cancer effects in vitro and in vivo. PLoS One. 2011;6.

73. Hellmich MR, Coletta C, Chao C, Szabo C. The Therapeutic Potential of Cystathionine β-Synthetase/Hydrogen Sulfide Inhibition in Cancer. Antioxid Redox Signal. 2015;22:424–48.

74. Richman S. Deficient mismatch repair: Read all about it (Review). International Journal of Oncology. 2015;47:1189–202.

75. Cai W, Wang M, Ju L, Wang C, Zhu Y. Hydrogen sulfide induces human colon cancer cell proliferation: role of Akt, ERK and p21. Cell Biol Int. 2010;34:565–72.

76. Purcell R V., Pearson J, Aitchison A, Dixon L, Frizelle FA, Keenan JI. Colonization with enterotoxigenic Bacteroides fragilis is associated with early-stage colorectal neoplasia. PLoS One. 2017;12.

77. Maiuri AR, Peng M, Sriramkumar S, Kamplain CM, DeStefano Shields CE, Sears CL, et al. Mismatch repair proteins initiate epigenetic alterations during inflammation-driven tumorigenesis. Cancer Res. 2017;77:3467–78.

78. Lupton JR. Microbial degradation products influence colon cancer risk: the butyrate controversy. J Nutr. 2004;134:479–82.

79. Magnúsdóttir S, Thiele I. Modeling metabolism of the human gut microbiome. Current Opinion in Biotechnology. 2018;51:90–6.

80. Benedict MN, Mundy MB, Henry CS, Chia N, Price ND. Likelihood-Based Gene Annotations for Gap Filling and Quality Assessment in Genome-Scale Metabolic Models. PLoS Comput Biol. 2014;10:e1003882. doi:10.1371/journal.pcbi.1003882.

81. Magnúsdóttir S, Heinken A, Kutt L, Ravcheev DA, Bauer E, Noronha A, et al. Generation of genome-scale metabolic reconstructions for 773 members of the human gut microbiota. Nat Biotechnol. 2016. doi:10.1038/nbt.3703.

